# Less Is More: A Mutation in the Chemical Defense Pathway of *Erysimum cheiranthoides* (Brassicaceae) Reduces Total Cardenolide Abundance but Increases Resistance to Insect Herbivores

**DOI:** 10.1101/2020.05.01.072850

**Authors:** Mahdieh Mirzaei, Tobias Züst, Amy P. Hastings, Anurag A. Agrawal, Georg Jander

## Abstract

Many plants produce structurally related defensive metabolites with the same target sites in insect herbivores. Two possible drivers of this chemical diversity are: (i) interacting effects of structurally related compounds increase resistance against individual herbivores, and (ii) variants of the same chemical structures differentially affect diverse herbivore species or feeding guilds. *Erysimum cheiranthoides* L (Brassicaceae; wormseed wallflower) produces abundant and diverse cardenolide toxins, which are derived from digitoxigenin, cannogenol, and strophanthidin, all of which inhibit Na^+^/K^+^-ATPases in animal cells. Here we describe an *E. cheiranthoides* mutant with 66% lower cardenolide content, resulting from greatly decreased cannogenol- and strophanthidin-derived cardenolides, partially compensated for by increases in digitoxigenin-derived cardenolides. This compositional change created a more even cardenolide distribution, decreased the average cardenolide polarity, but did not impact glucosinolates, a different class of chemical defenses. Growth of generalist herbivores from two feeding guilds, *Myzus persicae* Sulzer (Hemiptera: Aphididae; green peach aphid) and *Trichoplusia ni* Hübner (Lepidoptera: Noctuidae; cabbage looper), was decreased on the mutant line compared to wildtype. Both herbivores accumulated cardenolides in proportion to plant content, with *T. ni* accumulating higher total concentrations than *M. persicae.* Helveticoside, an abundant cardenolide in *E. cheiranthoides*, was absent in *M. persicae*, suggesting that this compound is not present in the phloem. Our results support the hypothesis that cardenolide diversity protects plants against different herbivores, with digitoxigenin-derived compounds providing better protection against insects like *M. persicae* and *T. ni*, whereas cannogenol and strophanthidin provide better protection against other herbivores of *E. cheiranthoides.*

**Funding:** This research was funded by US National Science Foundation awards 1907491 to AAA and 1645256 to GJ and AAA, Swiss National Science Foundation grant PZ00P3-161472 to TZ, and a Triad Foundation grant to GJ.

## INTRODUCTION

Plants produce a wide array of specialized metabolites as chemical defenses against insect herbivory. Although some individual compounds can be broadly effective, greater chemical diversity has been shown to provide enhanced protection against a variety of herbivores. For example, a detailed study of 31 sympatric tree species from the monophyletic clade Protieae (Burseraceae) showed that an increased diversity of chemical defenses, including procyanidins, flavone glycosides, chlorogenic acids, saponins, triterpenes and sesquiterpenes, was associated with enhanced herbivore resistance (Salazar et al. 2018).

Diversity of chemical structures, even if they belong to the same metabolic class, can provide defensive advantages. In some cases, compounds with similar structures and the same metabolic target site may act synergistically in defense against individual herbivore species. For instance, although they have comparable photosensitizing functions, six fouranocoumarins from *Pastinaca sativa* L (Apiaceae) provided enhanced defense against *Heliothis zea* (Lepidoptera: Noctuidae) relative to each compound individually (Berenbaum et al. 1991). Alternatively, a more varied chemical arsenal could be evolutionary advantageous if variants of the same chemical structure differ in their toxicity towards multiple herbivore species. In *Arabidopsis thaliana* L (Brassicaceae), which produces both tryptophan-derived indole glucosinolates and methionine-derived aliphatic glucosinolates, indole glucosinolates provide better protection against *Myzus persicae* Sulzer (Hemiptera: Aphididae; green peach aphid) (Kim et al. 2008). Conversely, growth of *Trichoplusia ni* Hübner (Lepidoptera: Noctuidae; cabbage looper), a generalist lepidopteran herbivore, was more strongly impacted by indole rather than aliphatic glucosinolates in *A. thaliana* mutant lines (Müller et al. 2010). Performance of the crucifer-feeding specialist *Pieris rapae* L (Lepidoptera: Pieridae; white cabbage butterfly), which has a high level of glucosinolate tolerance due to chemical diversion of the breakdown products (Wittstock et al. 2004), was not improved by the loss of either indole or aliphatic glucosinolates in *A. thaliana* (Müller et al. 2010).

*Erysimum cheiranthoides* L (Brassicaceae; wormseed wallflower), a widely distributed annual species, accumulates not only indole and aliphatic glucosinolates, but also cardenolides, steroid-derived specialized metabolites that function as inhibitors of essential Na^+^/K^+^-ATPases in animal cells (Züst et al. 2020). This additional set of defensive compounds provides enhanced protection against crucifer-specialist herbivores that can tolerate or detoxify glucosinolates, including the glucosinolate-tolerant *P. rapae* (Dimock et al. 1991; Renwick et al. 1989; Sachdev-Gupta et al. 1993; Sachdev-Gupta et al. 1990).

Cardenolides and structurally related bufadienolides, which also inhibit Na^+^/K^+^-in animal cells, are produced by at least a dozen plant families (Agrawal et al. 2012). Like glucosinolates, for which more than 120 different compounds have been identified across the Brassicaceae (Fahey et al. 2001), cardenolides are a large group of chemically and functionally similar plant defensive metabolites. A recent analysis identified a total of 95 different cardenolide structures in 48 species of the *Erysimum*, the only cardenolide-producing genus within the Brassicaceae (Züst et al. 2018; Züst et al. 2020). This vast diversity is primarily generated by modifications to the core steroid structure of the molecules (the genin), and by glycosylation of the genin with various sugar moieties to form linear glycoside ‘tails’. In *E. cheiranthoides*, the first *Erysimum* species to have its genome sequenced (Züst et al. 2020), the most abundant cardenolides are glycosides of the three genins digitoxigenin, cannogenol, and strophanthidin (Fig. 1).

**Fig. 1.**
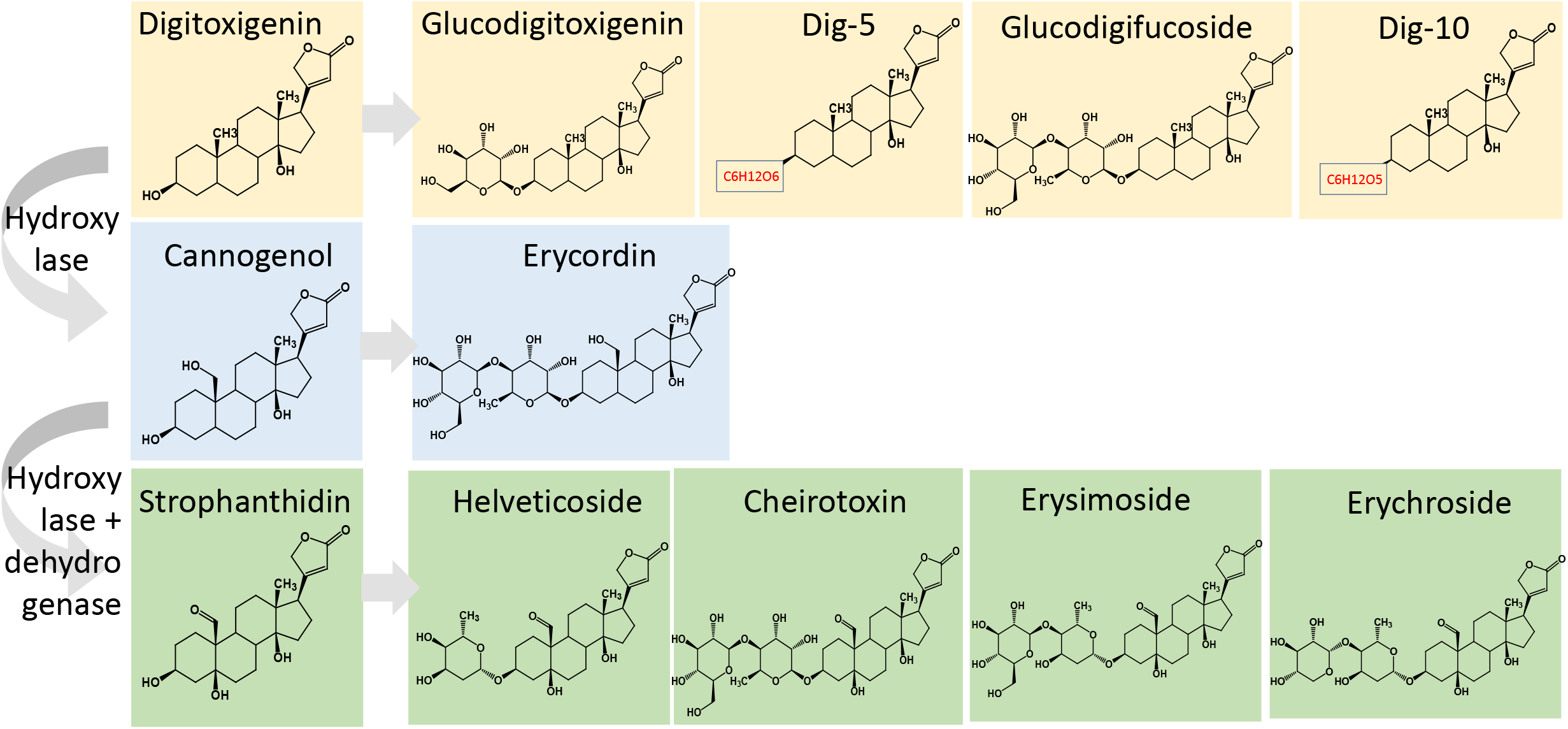
Structures of cardenolides found in *Erysimum cheiranthoides.* Predicted enzymes that would convert digitoxigenin to strophanthidin and then cannogenol are shown. Digitoxigenin-, cannogenol-, and strophanthidin-derived cardenolides are highlighted in yellow, blue, and green, respectively.

Several *in vitro* studies have demonstrated that modifications of the core steroid structure of cardenolides alter the inhibitory effects on mammalian Na^+^/K^+^-ATPases (Dzimiri et al. 1987; Farr et al. 2002; Paula et al. 2005; Schonfeld et al. 1985). Similarly, a collection of 17 cardenolides showed a wide range of activity against the Na^+^/K^+^-ATPases of a cardenolide-tolerant specialist herbivore, *Danaus plexippus* L (Lepidoptera: Nymphalidae), and two non-adapted species, *Euploea core* Cramer (Lepidoptera: Nymphalidae) and *Schistocerca gregaria* Forsskål (Orthoptera) (Petschenka et al. 2018). The relative order of potency of the tested cardenolides varied among the three tested insect species, suggesting that a more diverse cardenolide profile would optimize resistance to a wider array of herbivores.

*In vitro* enzyme inhibition assays provide only a partial view of the effects that cardenolides have when they are consumed by insect herbivores. For instance, efflux pumps and barriers to diffusion that prevent movement of cardenolides into the nervous system of some insect species (Groen et al. 2017; Petschenka et al. 2013) would not be relevant in an *in vitro* Na^+^/K^+^-ATPase inhibition assay. To initiate an investigation of the *in vivo* role of cardenolide diversity in defense against insect herbivores, we conducted an *E. cheiranthoides* mutant screen to identify plants with altered cardenolide profiles. Experiments with *T. ni* and *M. persicae*, broad generalist herbivores from two different feeding guilds, associated greater amounts of digitoxigenin-derived cardenolides, but overall lower cardenolide abundance, with enhanced herbivore resistance.

## METHODS AND MATERIALS

### Plant, Insects, and Growth Conditions

All experiments were conducted with the genome-sequenced *E. cheiranthoides* var. Elbtalaue (Züst et al. 2020), Arabidopsis Biological Resource Center (https://abrc.osu.edu) accession number CS29250. Plants were grown in Cornell mix (by weight 56% peat moss, 35% vermiculite, 4% lime, 4% Osmocote slow-release fertilizer [Scotts, Marysville, OH], and 1% Unimix [Scotts]) in Conviron (Winnipeg, CA) growth chambers with a 16:8 photoperiod, 180 μM m^−2^ s^−1^ photosynthetic photon flux density, 60% humidity, and constant 23 °C temperature.

*Trichoplusia ni* eggs were obtained from Benzon Research (www.benzonresearch.com) and were hatched on artificial diet (Southland Products, http://www.southlandproducts.net) in an incubator at 28 °C. A *P. rapae* colony, which was started with approximately 30 adults that were collected on the Cornell University campus in August, 2019, was maintained on *Brassica oleracea* L (Brassicaceae) cv. Wisconsin Golden Acre, under the same growth chamber conditions as those used for *E. cheiranthoides*. *Myzus persicae* were from a lab colony of a previously described “USDA” strain (Ramsey et al. 2014; Ramsey et al. 2007), which was maintained on *Nicotiana tabacum* L (Solanaceae) in a growth room with a 16:8 photoperiod and constant 23 °C temperature.

### Mutagenesis

Ethyl methanesulfonate (EMS) mutagenesis was performed as described previously for experiments with *A. thaliana* (Kim et al. 2006). Briefly, 10 g inbred line Elbtalaue seeds (approx. 48,000 seeds) were soaked with 40 ml of 100 mM pH 7.5 phosphate buffer in a 50 ml plastic tube and stored overnight at 4 °C. After decanting the buffer, 40 ml of fresh 100 mM phosphate buffer with 0.6% EMS (Sigma-Aldrich, St. Louis, MO) was added to the tube. The mixture was incubated at 23 °C for seven hours with gentle shaking. Seeds were rinsed 20 times with distilled water prior to planting in ten 25 × 50 cm nursery flats. After 15 weeks, M2 seeds were harvested as ten separate pools, one from each flat. Approximately 100 M2 seeds from each pool of M1 plants were planted in individual 100 cm^3^ pots. After four weeks, five 0.5 cm hole punches were taken from mature leaves as samples for cardenolide analysis from ~70 M2 plants grown from each of the 10 pools. Foliar cardenolides were analyzed by UPLC-MS as described below. Confirmed mutant plants were back-crossed to wildtype Elbtalaue and F2 progeny from these crosses were used for further experiments.

### Cardenolide Assays

Ground frozen leaf tissue, 50-100 mg per sample, was weighed into 2 ml tubes. After addition of four 3-mm stainless steel balls (Abbott Ball Company, Hartford, CT) to each tube, the samples were homogenized for 2 × 2 min at using a Harbil 5G-HD shaker (Fluid Management, Wheeling, IL) and kept frozen in liquid nitrogen until extraction. Three times tissue volume of methanol : formic acid (99.8% : 0.2%) were added. Samples were vigorously vortexed for 10 s, centrifuged for 30 s at 400 g, sonicated in an ultrasonic bath for 30 min, and then centrifuged for 30 s at 400 g. The extracts were centrifuged for 10 min at 20,000 g and the supernatants were transferred to glass HPLC vials for LCMS analysis using a Q-Exactive hybrid quadrupole-orbitrap mass spectrometer (Thermo Scientific) equipped with a Titan™ C18 UHPLC Column, (100 × 2.1 mm, particle size 1.9 μm; Sigma-Aldrich). 2-3 μl injections were separated using the following linear gradient: mobile phase B (100% acetonitrile plus 0.1% formic acid) increased from 2 to 97% over 10 min, held at 97% for 1.5 min, and returned to 98% mobile phase A (0.1% formic acid in water) for equilibration for 1.5 min. The flow rate of the mobile phases was 0.5 ml/min, and the column temperature and the autosampler temperature were maintained at 40 °C and 15 °C, respectively. Eluted compounds were detected from 200-800 m/z with 140,000 resolution (full width at half maximum, at m/z 200) in positive and negative ion modes.

Cardenolide compounds were relatively quantified by isolating the following positive-ion chromatograms: cheirotoxin (C35H52O15), rt = 3.93 min, [M+H]^+^= 713.3384 m/z; erysimoside (C35H52O14), rt = 4.06 min, [M+H]^+^= 697.3435 m/z; erychroside (C34H50O13), rt = 4.2 min, [M+H]^+^= 667.3329 m/z; helveticoside (C29H42O9), rt = 4.65 min, [M+H]^+^= 535.2907 m/z,; digitoxigenin (C23H34O4), rt = 5.8 min, [M+H]^+^= 375.2535 m/z; glucodigitoxigenin (C29H44O9), rt = 4.46 min, [M+H]^+^= 537.3063 m/z; dig-5 (C29H44O9), rt = 4.55 min, [M+H]^+^= 537.3063 m/z; glucodigifucoside (C35H54O13), rt = 4.7 min, [M+H]^+^= 683.3642 m/z; dig-10 (C29H44O8), rt = 5.09 min, [M+H]^+^= 521.3114 m/z; erycordin (C35H54O14), rt = 4.15 min, [M+H]^+^= 699.3592 m/z. In assays of insect samples, the sodium adducts of two cardenolides, cheirotoxin [M+Na]^+^= 735.3187 m/z and helveticoside [M+Na]^+^= 557.2721 m/z, were the most abundant and were used for relative quantification. Glucosinolate compounds were relatively quantified by isolating the following negative-ion chromatograms: indol-3-ylmethyl glucosinolate (C16H20N2O9S2; I3M), rt = 1.85 min, [M-H]^−^= 447.0532 m/z; 4-methoxy-indol-3-ylmethyl glucosinolate (C17H22N2O10S2; 4MOI3M), rt = 2.36 min, [M-H]^−^= 477.0638 m/z; 4-hydroxy-indol-3-ylmethylglucosinolate (C16H20N2O10S2; 4OHI3M), rt = 0.9 min, [M-H]^−^= 463.0484 m/z; 3-methylsulfinylpropyl glucosinolate (C11H21NO10S3; 3MSIP), rt = 0.56 min, [M-H]^−^= 422.0249 m/z; 3-methylsulfonylpropyl glucosinolate (C11H21NO11S3; 3MSOP), rt = 0.55 min, [M-H]^−^= 438.0196 m/z; 4-methylsulfonylbutyl glucosinolate (C12H23NO11S3; 4MSOP), rt = 0.58 min, [M-H]^−^= 452.0360 m/z; N-methylbutyl glucosinolate (C12H23NO9S2; NMB), rt = 1.8 min, [M-H]^−^= 388.0739 m/z.

Digitoxigenin (Sigma-Aldrich), K-strophanthin (a mixture of cymarin, erysimoside, helveticoside, strophanthidin, k-β-strophanthin, and k-strophanthoside; Sigma-Aldrich), and helveticoside (Cfm Oskar Tropitzsch GmbH, Germany) standards were purchased commercially, and the remaining compounds were validated using ms/ms analysis and comparison to known *E. cheiranthoides* cardenolide structures. The resulting raw LC/MS files were uploaded into XCMS online (https://xcmsonline.scripps.edu/) and processed for untargeted metabolomics profiling using predefined parameter sets for the UPLC/Q-Exactive system. The normalized targeted mass features corresponding to cardenolides and glucosinolates were then isolated from the output file.

An HPLC-UV assay was performed to quantify the absolute amount of cardenolides in both wildtype and mutated plant samples with seven biological replicates. Ten mg freeze-dried ground plant tissue was weighed into a screw cap tube with ~ 30 zirconia/silica beads (2.3 mm; BioSpec Products, Bartlesville, OK). One ml digitoxin standard solution (19.61 μg digitoxin dissolved in 1 ml HPLC-grade 100% methanol) was added to each tube and the samples were immediately vortexed for 10 s, shaken on a FastPrep shaker (MP Biomedicals, Irvine, CA) twice for 45 s at 6.5 m/s, and centrifuged for 12 min at 21,000 g. The supernatants were transferred to new labeled 1.5 ml tubes. The supernatants were transferred to new labeled 1.5 ml tubes, and the extracts dried down completely in a Centrivap (Labconco, Kansas City, MO) at 30 °C, reconstituted in 100 μl absolute methanol and vortexed for approximately 30 s to dissolve the dried residue. The samples were then transferred into a syringe filter (Kinesis 13 mm, 0.45 μm; Cole-Parmer, East Bunker, CT), and pushed through into an HPLC vial with an insert (300 μl, pulled-point). Fifteen μl of the extract were injected into an Agilent 1100 series HPLC and compounds were separated on a Gemini C18 reversed phase column (5 μm, 250 mm × 10 mm, Macherey-Nagel, Düren, Germany). Cardenolides were eluted with a constant flow of 0.7 ml/min using the following gradient: 0–2 min 16% acetonitrile, 25 min 70% acetonitrile, 30-40 min 95% acetonitrile, and a 10 min post-run at 16% acetonitrile. UV absorbance spectra were recorded at 218 nm using a diode array detector (Agilent Technologies, Santa Clara, CA). Erysimoside, helveticoside, digitoxigenin and digitoxin peaks were validated using commercially available standards that were run separately from the experimental samples. The other cardenolides were identified based on the order of elution in LCMS experiments as the following retention times: cheirotoxin, rt = 9.6 min; erysimoside, rt = 10 min; erychroside, rt = 10.9 min; helveticoside, rt = 12.3 min; digitoxigenin, rt = 16.6 min; glucodigitoxigenin, rt = 12 min; dig-5, rt = 13.1 min; glucodigifucoside, rt = 13.3 min; dig-10, rt = 14.4 min; erycordin, rt = 10.4 min, and digitoxin, rt = 19 min. The individual total cardenolide content was calculated based on the peak areas relative to the internal digitoxin standard (19.6 μg/ml) of each sample.

### Insect Bioassays

Three days after hatching, *T. ni* larvae were placed individually on leaves of F2 progeny of EMS-mutagenized plants backcrossed to Elbtalaue. Larvae were confined on individual *E. cheiranthoides* leaves using 6.5 × 8 cm organza bags (Supplemental Fig. 1A; www.amazon.com, item B073J4RS9C). After ten days, the larvae were harvested, weighed individually, and frozen immediately in liquid nitrogen. Experiments with *P. rapae* were conducted in the same manner, but larvae were not weighed because they did not survive.

Aphids were caged on *E. cheiranthoides* leaves in groups of 5 fourth-instar aphids. Cages were prepared from cut 50-ml plastic tubes (segments of 3 cm height and 3 cm diam) with a fine-mesh covered lid (Supplemental Fig. 1B). After 14 days, the surviving adult aphids and nymphs were counted and frozen immediately in liquid nitrogen. Frozen caterpillar and aphid samples (10-20 mg each) were extracted and analyzed by HPLC-MS, as described above for plant samples above.

As bodies of aphids feeding in *E. cheiranthoides* were devoid of helveticoside, uptake of this cardenolide was tested in an artificial diet assay. Aphid artificial diet (Douglas et al. 2001) consisted of 500 mM sucrose, 150 mM amino acids (Ala, 5.7 mM; Arg, 14.3 mM; Asn, 14.3 mM; Asp, 14.3 mM; Cys, 2.7 mM; Glu, 8.4 mM; Gln, 16.5 mM; Gly, 1.2 mM; His, 8.7 mM, Ile; 8.7 mM; Leu, 8.7 mM; Lys, 8.7 mM; Met, 2.9 mM; Phe, 2.9 mM; Pro, 5.7 mM; Ser, 5.7 mM; Thr, 8.7 mM; Trp, 2.9 mM; Tyr, 0.6 mM; Val, 8.7 mM), and a mixture of essential minerals and vitamins (final diet pH 7-7.5). Helveticoside was first dissolved in absolute ethanol to make a 0.05 mg/μl stock solution and then a 100 ng/μl working solution was prepared by adding from the stock solution to the fresh diet. The diet was prepared at different concentration of helveticoside (0, 20, 50 ng/μl). A quantity of 600 μl filter-sterilized (0.2 μm) diet was placed between two stretched layers of parafilm over the edge of the above-described cages. Forty to sixty adult aphids were placed on the underside of each diet cage and were allowed to feed for three days and then frozen immediately in liquid nitrogen for the further HPLC-MS analysis.

### Na^+^/K^+^-ATPase Inhibition Assay

We measured porcine (*Sus scrofa* L) Na^+^/K^+^-ATPase activity in the presence of plant extracts, following the protocol in Züst et al. (2019), to test the inhibitory effect of the cardenolides in wildtype and 454 mutant *E. cheiranthoides* samples. To prepare sample extracts, 10 mg freeze-dried ground plant tissue was weighed into a screw cap tube with ~ 30 zirconia/silica beads (2.3 mm; BioSpec Products, Bartlesville, OK). One ml absolute methanol was added to each tube and tubes were shaken on a FastPrep shaker (MP Biomedicals, Irvine, CA) twice for 45 s at 6.5 m/s and centrifuged for 10 min at 21,000 g. Seven hundred μl supernatants were transferred to new labeled 1.5 ml microcentrifuge tubes. The extracts were dried down completely in a Centrivap (Labconco, Kansas City, MO) at 30 °C, and resuspended in 50 μL 100% dimethyl sulphoxide (DMSO) by vortexing for 20 s, sonicating in an ultrasonic bath for 2 × 5 min, and briefly centrifuging (10 s) at 16,0000 g. Extracts were diluted 10-fold with deionized water (EMD Millipore, Burlington, MA) to reach a concentration of 10% DMSO. A series of six dilutions (0.3, 0.09, 0.027, 0.0081, 0.0024, 0.00073) was prepared from each full-strength sample extract, using 10% DMSO. Six biological replicates of wildtype samples and seven biological replicates of mutated samples were distributed randomly between two 96-well plates.

Enzyme assays were carried out as in Züst et al. (2019), with ouabain standards (final concentration of 10^−3^ to 10^−8^ M, in 2% DMSO) included on each plate for reference. A replicated set of the dilution series, with the same reaction conditions but excluding KCl, was included for each plant sample. ATPase activity was quantified as percent control activity, by first subtracting from the reaction absorbance its appropriate background (absorbance of equivalent reaction without KCl), and then by comparison to an uninhibited control. For each plant sample, a logistic curve was fit to the percent enzyme activity data, as in Züst et al. (2019), except that here we used a 4-parameter function to estimate the upper and lower asymptote for each curve. Cardenolide concentrations of the *E. cheiranthoides* leaf samples were then calculated in terms of ouabain equivalents.

### Data Analysis

Statistical comparisons were made using Microsoft Excel and JMP v. 10 (www.jmp.com). The evenness index of the cardenolide distribution was calculated using the Evar method (Smith and Wilson 1996). The polarity index of mutant and wildtype *E. cheiranthoides* cardenolides was calculated by multiplying the proportion of each cardenolide of the total by its retention time in the HPLC-UV assay, and averaging this value across all cardenolides in the sample (Rasmann and Agrawal 2011).

## RESULTS

In an initial screen of 625 EMS-mutagenized *E. cheiranthoides* M2 plants by UPLC-MS, we were able to reliably quantify the abundance of eight known cardenolides: digitoxigenin, two digitoxigenin-derived glycosides (glucodigitoxigenin and glucodigifucoside), a cannogenol-derived glycoside (erycordin), and four strophanthidin-derived glycosides (cheirotoxin, erysimoside, erychroside, and helveticoside). Additionally, we observed two likely cardenolides with a genin fragmentation pattern consistent with digitoxigenin (labeled Dig-5 and Dig-10, respectively). These compounds have molecular weights of 537 and 521, respectively, suggesting glycosylation with glucose for Dig-5 and deoxyhexose for Dig-10, although the exact structures have not been determined.

Plants with divergent phenotypes for one or more cardenolides (Fig. 2A; with glucodigifucoside as an example) were identified and subjected to further screening that resulted in five confirmed mutant lines. Mutant line 454 accumulated elevated digitoxigenin-derived cardenolides and low concentrations of cannogenol- and strophanthidin-derived cardenolides (Fig. 2B). Mutants 634, 812, 826, and 894 accumulated significantly decreased amounts of most or all cardenolides (Fig. S2A-D). Two of these mutants, 812 and 894, originated from the same M1 seed pool and may be siblings.

**Fig. 2.**
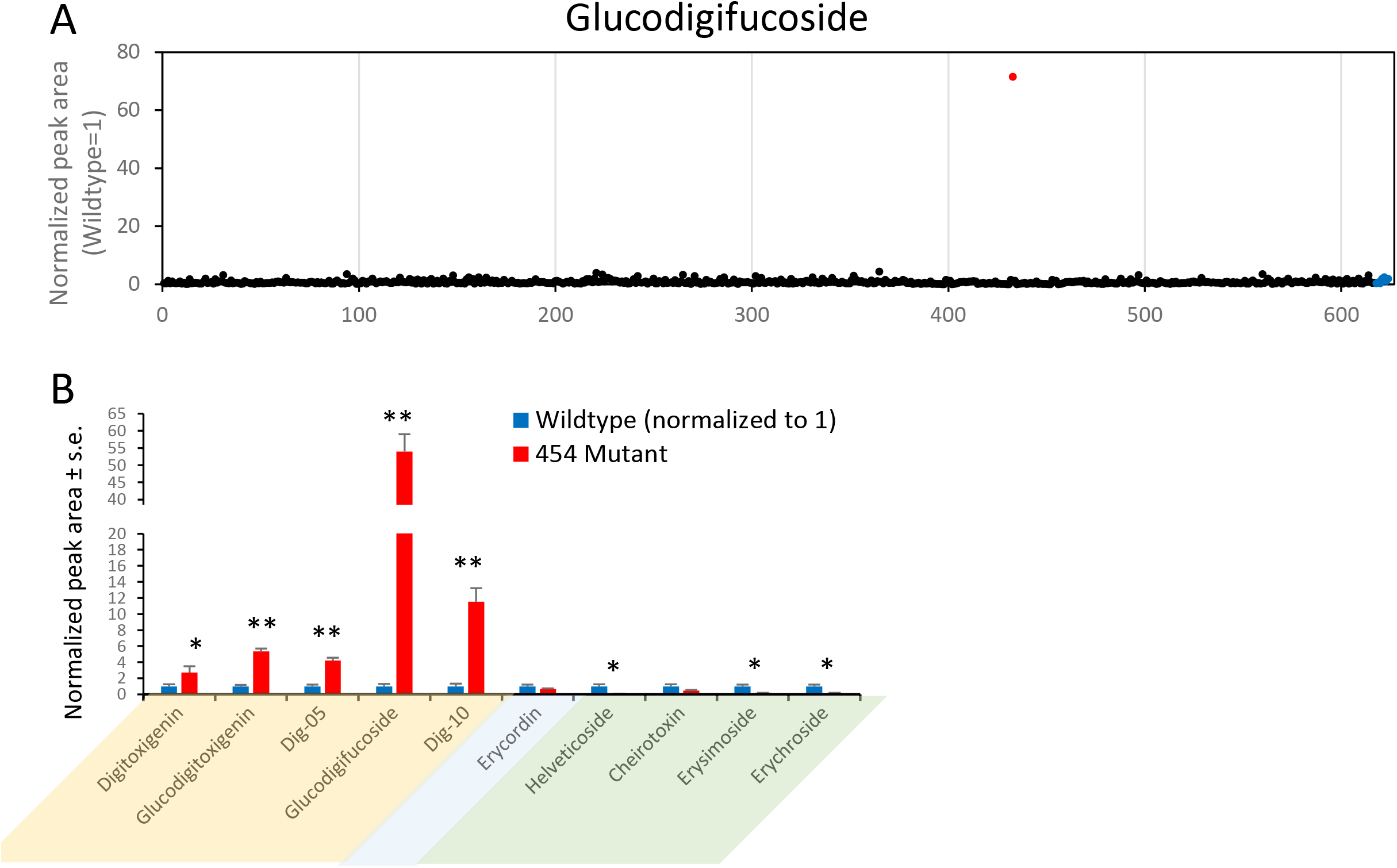
Mutant screen for altered cardenolide content. (A) Glucodigifucoside abundance at the first screening of 625 M2 mutant lines. Each dot represents a single plant measured by UPLC-MS. The high outlier at center-right represents the 454 mutant line and the rightmost seven dots represent wildtype Elbtalaue samples. (B) Cardenolide profile of mutant line 454, as measured by UPLC-MS, with wildtype normalize to 1 for each cardenolide. Mean ± s.e. of N = 5, *P < 0.05, **P < 0.01, t-test.

The 454 mutant, which had the most distinctive phenotype, was crossed to wildtype Elbtalaue and cardenolides were measured in 203 F2 progeny. Among these progeny, 155 had a wildtype cardenolide phenotype and 48 had the 454 mutant phenotype (Fig. S3), a roughly 3:1 segregation pattern that indicated a single recessive mutation. Relative to their wildtype siblings, the 454 mutant plants had elevated digitoxigenin-derived cardenolides and decreased cannogenol- and strophanthidin-derived cardenolides (Fig. S3), similar to the original 454 mutant line.

As a more quantitative assay of the cardenolide abundance in the 454 mutant and phenotypically wildtype siblings, we used HPLC-diode array detection at 218 nm, which measures absorbance of the lactone ring that is common to all of the cardenolide structures. In particular, erychroside and other strophanthidin-derived cardenolides are poorly detected in the UPLC-MS assay (Fig. 2B) relative to the UV absorption assay (Fig. 3A). Nevertheless, similar to the MS detection, the 218 nm absorption assay showed increased digitoxigenin-derived cardenolides and decreased cannogenol- and strophanthidin-derived cardenolides (Fig. 3A). Notably, the 454 mutation caused a 92% decrease in erychroside. The total cardenolide UV absorbance was decreased by 66% in the 454 mutant relative to wildtype Elbtalaue (Fig. 3B). Although the same cardenolides were present in mutant and wildtype plants, the evenness of the cardenolide distribution was higher in the mutant (Fig. 3C). An index of compound polarity, calculated as HPLC retention time multiplied by relative abundance of each cardenolide (Rasmann and Agrawal 2011), was substantially higher in the mutant than in wildtype, indicating that the mutant cardenolide profile is less polar (Fig. 3D). With the exception of N-methylbutyl glucosinolate, which was slightly increased in the 454 mutant line, there were no significant glucosinolate changes relative to wildtype (Fig. 3E).

**Fig. 3.**
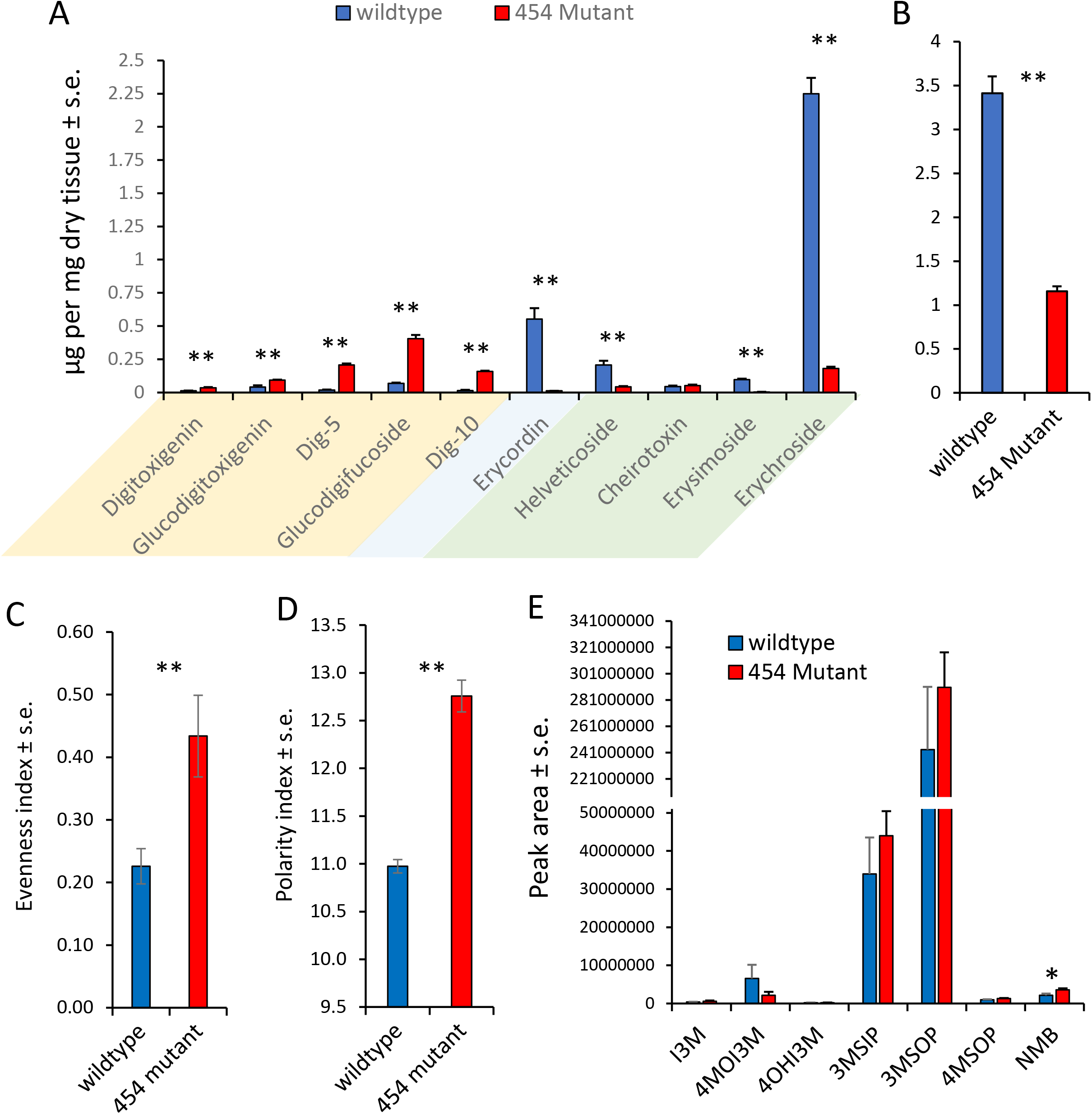
Cardenolide and glucosinolate content of 454 mutant line relative to wildtype Elbtalaue. (A) Abundance of each individual cardenolide, and (B) total amount measured by HPLC-UV. Digitoxigenin-, cannogenol-, and strophanthidin-derived cardiac glycosides are highlighted in yellow, blue, and green, respectively. (C) Evenness of the cardenolide distribution. (D) Polarity index, (E) Relative glucosinolate content of 454 mutant line compare to wildtype measured by LCMS. Mean ± s.e. of N = 7, *P < 0.05, **P < 0.01, *t*-test comparing wildtype and mutant. I3M = indol-3-ylmethyl glucosinolate, 4MOI3M = 4-methoxy-indol-3-ylmethyl glucosinolate, 4OHI3M = 4-hydroxy-indol-3-ylmethylglucosinolate, 3MSIP = 3-methylsulfinylpropyl glucosinolate, 3MSOP = 3-methylsulfonylpropyl glucosinolate, 4MSOP = 4-methylsulfonylbutyl glucosinolate, NMB = N-methylbutyl glucosinolate

We used an assay based on the inhibition of pig brain Na^+^/K^+^-ATPase (Petschenka et al. 2018) to determine whether the 454 mutant line has altered inhibitory activity relative to its wildtype Elbtalaue progenitor. Consistent with the greater abundance of cardenolides in the wildtype samples (Fig. 3B), extracts of wildtype Elbtalaue inhibited Na^+^/K^+^-ATPase activity to a greater extent than extracts of the 454 mutant (Fig. 4A). Based on comparisons to inhibition by a dilution series of ouabain, the 454 mutant line was calculated to have a 59% decrease in Na^+^/K^+^-ATPase activity compared to wildtype Elbtalaue (Fig. 4B).

**Fig 4.**
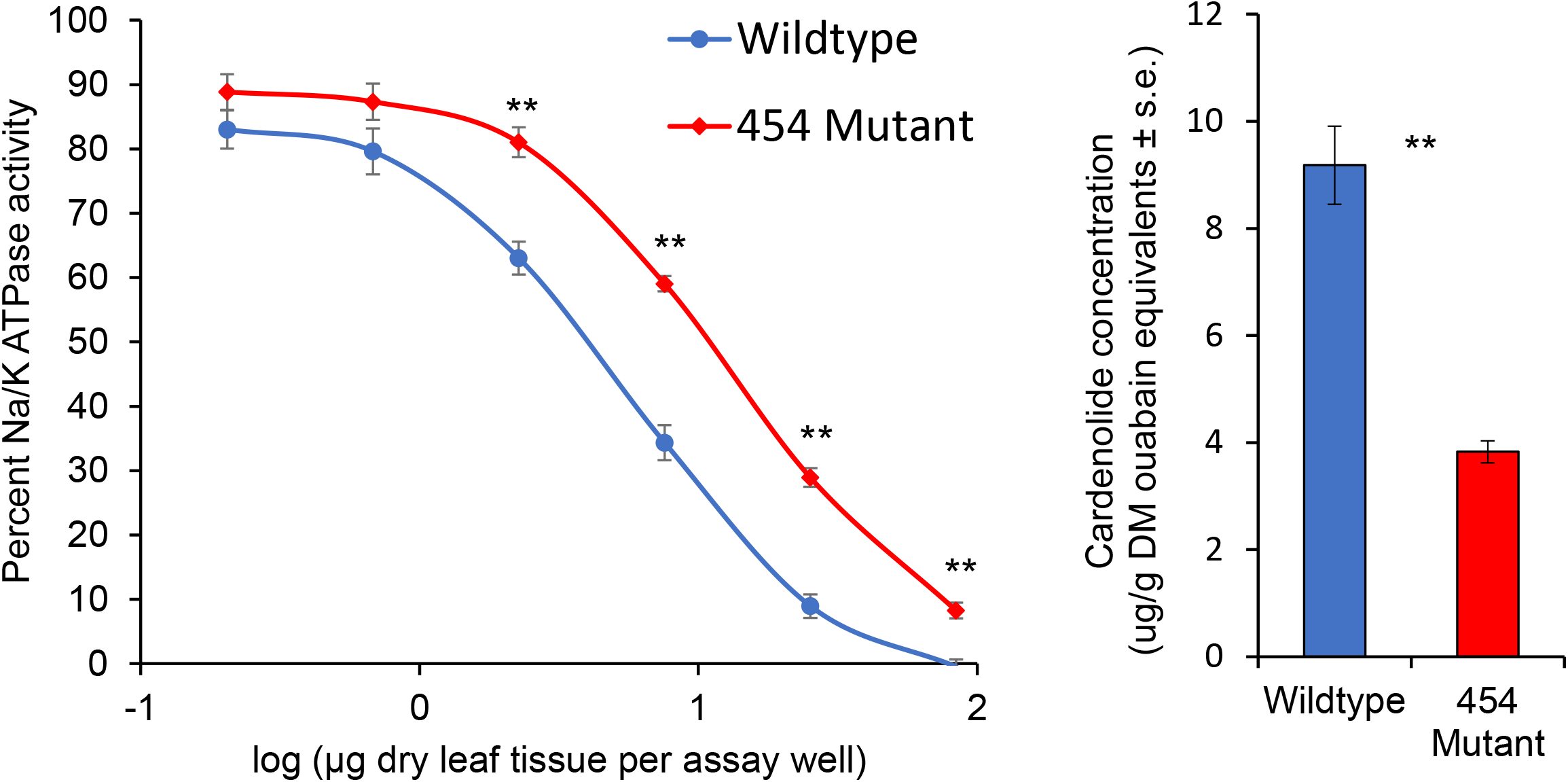
Inhibition of pig brain Na+/K+ ATPase. (A) Percent inhibition by dilutions of extracts of wildtype Elbtalaue and 454 mutant *E. cheiranthoides.* (B) Amount of inhibitory activity expressed as an equivalent amount of ouabain. *\**\*P < 0.01, t-test, mean ± s.e. of N = 7.

The segregating F2 population was used for bioassays with three insect herbivores, *T. ni*, *M. persicae*, and *P. rapae*. Relative to plants with wildtype cardenolide content, *T. ni* larvae on 454 mutant plants were smaller (Fig. 5A,B). Survival of adult *M. persicae* was not affected by host plant, but aphids produced 54% fewer progeny on the 454 mutant than on wildtype Elbtalaue (Fig. 5C). Crucifer-specialist *P. rapae* larvae, which are highly sensitive to cardenolides (Sachdev-Gupta et al. 1993), did not survive on either mutant or wildtype plants.

**Fig. 5.**
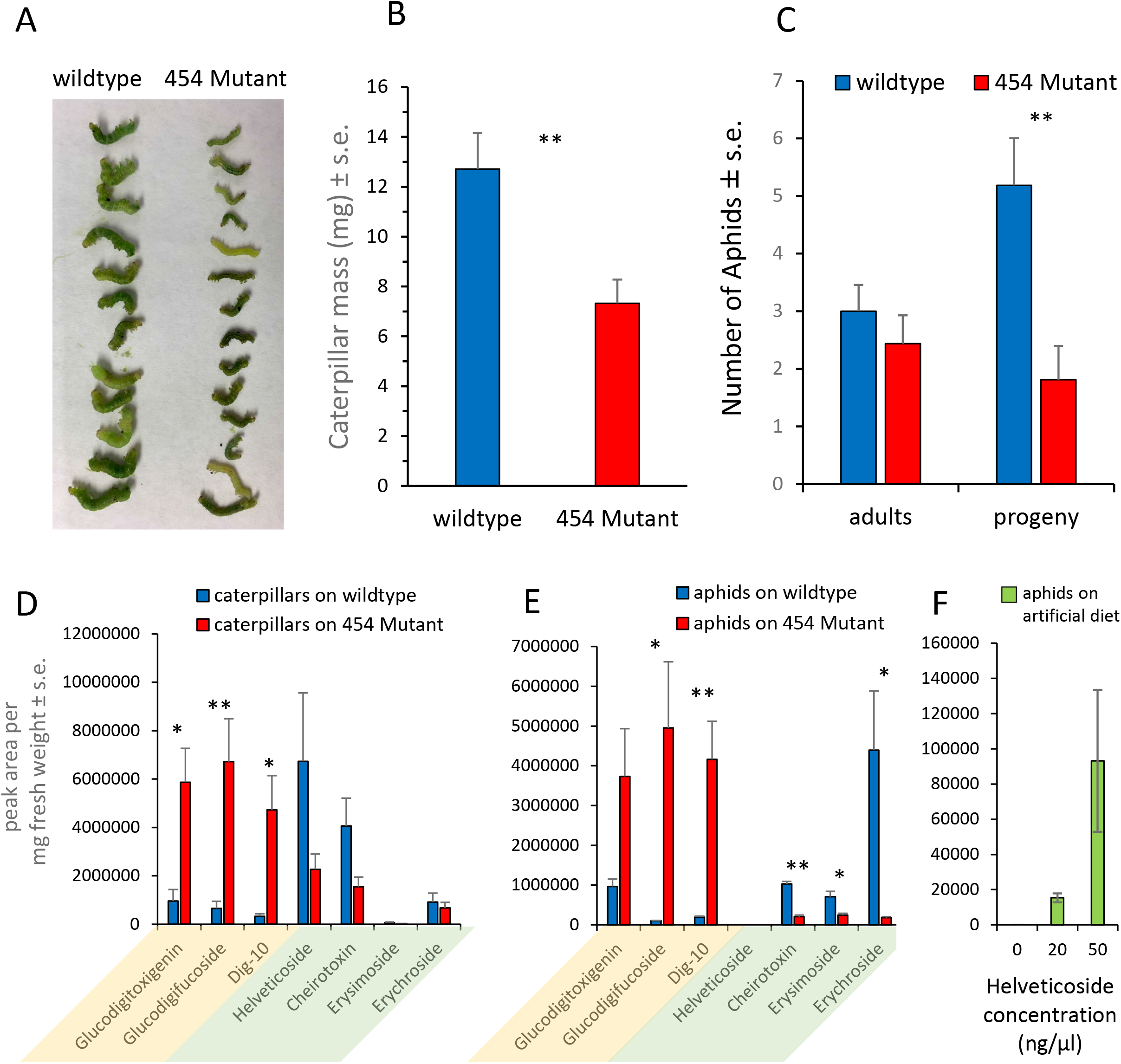
Insect growth on 454 mutant plants and segregating wildtype progeny from a backcross to wildtype Elbtalaue. (A) Photograph of a subset of the *Trichoplusia ni* larvae after 10 days of feeding. (B) *T. ni* larval mass, Mean +/-s.e. of N = 37. (C) Number of surviving adult aphids and progeny after two weeks feeding. Mean +/-s.e. of N = 16. Cardiac glycoside content in (D) *T. ni* caterpillars and (E) *M. persicae* feeding on sibling plants with wildtype and 454 mutant cardenolide content. Mean ± s.e. of N = 5. *P < 0.05, **P < 0.01, *t-*test. (F) Helveticoside content in *M. persicae* feeding from artificial diet with helveticoside. Mean +/-s.e. of N = 5.

Both *T. ni* and *M. persicae* accumulated cardenolides in their bodies as they were feeding, with cardenolides that were more abundant in the host plants (Fig. 3A) generally also accumulating to a higher level in the insects (Fig. 5D,E). Dig-5, digitoxigenin, and erycordin were not reliably detected in the insect bodies and were therefore not quantified. Most of the measured cardenolides were more abundant in *T. ni* larvae than in *M. persicae* (Table 1). However, erysimoside, as well as erychroside when the insects were feeding on wildtype Elbtalaue, were more abundant in aphids than in caterpillars. Helveticoside was not detected in aphids feeding on either wildtype or mutant *E. cheiranthoides* (Fig. 5E). However, when helveticoside was added to aphid artificial diet at a concentration similar to that found in *E. cheiranthoides* leaves, helveticoside was detected in the aphid bodies by UPLC-MS (Fig. 5F).

**Table 1.** Cardenolide abundance in insects feeding on *E. cheiranthoides*, peak area/mg in *M. persicae* relative to *T. ni*

|  | *M. persicae*/*T. ni* ratio |  |
| --- | --- | --- |
|  | Wildtype | 454 Mutant |
| Glucodigitoxigenin | 1.00 | 0.64 |
| Glucodigifucoside | 0.16 | 0.74 |
| Dig-10 | 0.59 | 0.88 |
| Erycordin | 0.23 | 0.21 |
| Helveticoside | 0.00 | 0.00 |
| Cheirotoxin | 0.25 | 0.13 |
| Erysimoside | 11.34 | 10.79 |
| Erychroside | 4.80 | 0.27 |

## DISCUSSION

The molecular characterization of mutations and natural variation can facilitate research on the defensive functions of plant specialized metabolites, which are often produced as classes of compounds with similar chemical structures and target sites in insect herbivores. Until fairly recently, such experiments were only feasible with classes of defensive metabolites that are present in well-studied model systems, such as glucosinolates in *A. thaliana*, benzoxazinoids in *Zea mays* L (Poaceae), and glycoalkaloids in *Solanum lycopersicum* L (Solanaceae). However, advances in genome sequencing and molecular methods have made it feasible to initiate such research using non-model systems such as cardenolide production in *E. cheiranthoides.*

A screen of mutagenized *E. cheiranthoides* var. Elbtalaue identified mutant line 454, which had a dramatic shift in its cardenolide profile. This shift resulted in a 66% decrease in total cardenolide content that was due to lower accumulation of cannogenol- and strophanthidin-derived cardenolides, which was only partially compensated for by an increase in digitoxigenin-derived cardenolide (Fig. 3B). Consistent with the decrease in total cardenolide content, extracts of the 454 mutant inhibited the highly sensitive porcine Na^+^/K^+^-ATPase activity 59% less than extracts of wildtype Elbtalaue (Fig. 4), indicating that the different *E. cheiranthoides* cardenolides do not fundamentally differ in their inhibitory activity in this *in vitro* enzyme assay. This observation is consistent with a survey of 28 *Erysimum* species, where despite a vast diversity in cardenolide compounds, the total cardenolide HPLC-MS peak area was strongly correlated with Na^+^/K^+^-ATPase inhibition (r = 0.95, p<0.001; Züst et al. 2020). The shift in cardenolide profile of the 454 mutant line also resulted in a more even cardenolide distribution (Fig. 3C), which was largely due to a greater than 90% decrease in the amount of erychroside, a strophanthidin-derived polar cardenolide that is the most abundant cardenolide in wildtype *E. cheiranthoides*.

The 3:1 genetic segregation of the 454 mutant cardenolide profile in the F2 generation (Fig. S2) indicates that this phenotype is caused by a single recessive mutation. Structural comparisons suggest that digitoxigenin, likely the most basal of the three genins, would be converted to cannogenol by a hydroxylase, possibly a cytochrome P450 (Fig. 1). The cannogenol and strophanthidin classes of cardenolides could be eliminated by a knockout mutation of such an enzyme. However, it is also possible that a regulatory mutation reduces conversion of digitoxigenin to cannogenol and strophanthidin. For instance, in *A. thaliana* knockout mutations in two cytochrome P450 genes, *CYP79B* and *CY79B3*, eliminate indole glucosinolate production (Zhao et al. 2002), whereas mutations in two transcription factor genes, *MYB28* and *MYB29*, eliminate aliphatic glucosinolate production (Hirai et al. 2007). Map-based cloning of the 454 mutation will be required to determine whether a biosynthetic or regulatory gene of the cardenolide pathway in *E. cheiranthoides* is affected. Other *E. cheiranthoides* mutations, which caused more uniform decreases in total cardenolide content (Fig. S2A-D), may affect earlier steps in the biosynthetic pathway and are worthwhile targets for future studies.

Despite a 66% reduction in total cardenolide content, both *T. ni* and *M. persicae* grew less well on the 454 mutant line than on wildtype Elbtalaue (Fig. 5A-C). On both mutant and wildtype *E. cheiranthoides*, the *T. ni* larvae were smaller and *M. persicae* produced fewer progeny than expected for a more preferred host such as *A. thaliana* under similar growth conditions (Casteel et al. 2014; Jander et al. 2001; Kim and Jander 2007; Müller et al. 2010). Since *A. thaliana* and *E. cheiranthoides* have comparable amounts of glucosinolates and glucosinolate levels were largely unchanged in the 454 mutant line relative to the wildtype, the changes in herbivore growth are unlikely due to the glucosinolate defense. We cannot rule out the possibility that the small increase in the structurally uncharacterized N-methylbutyl glucosinolate in the mutant line (Fig. 3E) may have contributed to enhanced insect resistance in the 454 mutant. However, as *M. persicae* is impervious to methionine-derived aliphatic glucosinolates (Kim et al. 2008) and a total loss of aliphatic glucosinolates in *A. thaliana* is required to cause a 50% increase in *T. ni* growth (Müller et al. 2010), it is more likely that changes in cardenolides were the main cause of decreased herbivore growth.

Three scenarios may account for the reduced insect growth on the 454 mutant line despite the lower total amounts of cardenolides in these plants (Fig. 4): (i) in contrast to the pig enzyme, insect enzymes may be more sensitive to digitoxigenin-derived cardenolides; (ii) there is differential uptake of cardenolides in insects; or (iii) the two insect species can detoxify or excrete polar cardenolides more easily than apolar cardenolides. Although we currently lack information on the specific Na^+^/K^+^-ATPase sensitivities of *T. ni* and *M. persicae*, experiments with pig, *Drosophila*, and non-specialist lepidopteran species showed similar cardenolide sensitivities (Karageorgi et al. 2019), making the first scenario perhaps less likely. In contrast, previous research with *M. persicae* and three other aphid species feeding on *Asclepias* spp. (milkweed) showed that the aphids accumulated predominantly apolar cardenolides in their bodies while they could excrete polar cardenolides with their honeydew (Züst and Agrawal 2015), suggesting that uptake and excretion may play an important role in our study as well., Erychroside, most abundant cardenolide in wildtype *E. cheiranthoides*, is the third-most polar cardenolide produced by this species and, despite its high abundance, both *T. ni* and *M. persicae* accumulated this compound at markedly lower proportions relative to other cardenolides. This suggests that both herbivores were able to partially detoxify or excrete this compound. In *T. ni*, this detoxification may have involved cleaving of the outer xylose sugar, resulting in an increased accumulation of helveticoside (Fig. 5D), which was found to have no deterrent effects on the otherwise highly cardenolide-susceptible *P. rapae* (Renwick et al. 1989; Sachdev-Gupta et al. 1990). Aphids may have been able to rapidly excrete erychroside without ever taking it up into their body cavity, although further evaluation of cardenolide profiles in aphid honeydew will be necessary to confirm this. Assuming that the most abundant cardenolide compound of wildtype plants is largely ineffective against these insect herbivores, the shift to a higher proportion of digitoxigenin cardenolides and a more apolar profile in the 454 mutant line may therefore have increased the cardenolide load of both herbivores despite lower total amounts in the plant. This increased cardenolide load could then have caused a reduction in performance due to direct toxicity or by requiring larger investment into general detoxification mechanisms.

Differential accumulation of *E. cheiranthoides* cardenolides by the two insect species (Table 1, Fig. 5D-E) may at least in part be reflective of their feeding styles. Whereas *T. ni* larvae consume leaf tissue, *M. persicae* feed exclusively from the phloem. Higher accumulation of erysimoside by *M. persicae* may indicate that this cardenolide is relatively more abundant in the phloem than in the leaf as a whole. Conversely, the complete absence of helveticoside in aphids feeding from both wildtype and mutant *E. cheiranthoides* suggests that this cardenolide may be low or absent in the phloem. When helveticoside was added to aphid artificial diet at concentrations comparable to those found in *E. cheiranthoides* leaves (>20 ng/mg wet weight in wildtype Elbtalaue), helveticoside could be detected in the aphid bodies (Fig. 5F). Therefore, the absence of helveticoside in *M. persicae* feeding on *E. cheiranthoides* is likely not due to degradation of helveticoside by the aphids or inhibited uptake from the aphid gut. Helveticoside is unique among *E. cheiranthoides* cardenolides as its glycoside moiety consists solely of a digitoxose sugar (Fig. 1). In a large comparison of cardenolide uptake by *Digitalis lanata* cells, Christmann et al. (1993) concluded that cardenolides with terminal glucose sugars are phloem mobile, whereas those with terminal digitoxose sugars are not. Thus, while our observations suggest that at least xylose as terminal sugar (for erychroside) can enable phloem mobility of cardenolides as well, they support a complete lack of mobility for helveticoside.

The 454 mutant line was more resistant against insect herbivores from two different feeding guilds than the wildtype plant, despite producing an overall lower quantity of cardenolide defenses. At first glance, it thus appears that the 454 mutant line should represent a superior evolutionary strategy that should quickly outcompete the wildtype if it ever arose naturally. However, to fully understand the evolutionary drivers of these defense phenotypes, it will be essential to evaluate plant performance in the native habitat of *E. cheiranthoides*, as we may well be missing important constraints or antagonists that favor the wildtype phenotype. For example, larger mammalian herbivores may respond more to the bitter taste of cardenolides rather than their toxicity, in which case it may be more beneficial for plants to produce large quantities of a putatively ‘cheaper’ compound. Yet even without knowing all evolutionary drivers of the *E. cheiranthoides* defense phenotype, we can nonetheless conclude that a more diverse mixture of cardenolides serves to cope with the diversity of antagonists that threaten this plant.

Our results demonstrate the feasibility of using a mutagenesis approach to study the *in vivo* function of cardenolides in defense against insect herbivores. Isolation of additional mutations affecting the accumulation of individual cardenolides will enable research on their *in vivo* functions in plant defense. Moreover, genetic mapping of such mutations will make it possible to identify genes in the cardenolide biosynthetic pathway, which remains mostly unidentified in *E. cheiranthoides* and other cardenolide-producing plant species.

---

**Fig. S1.**
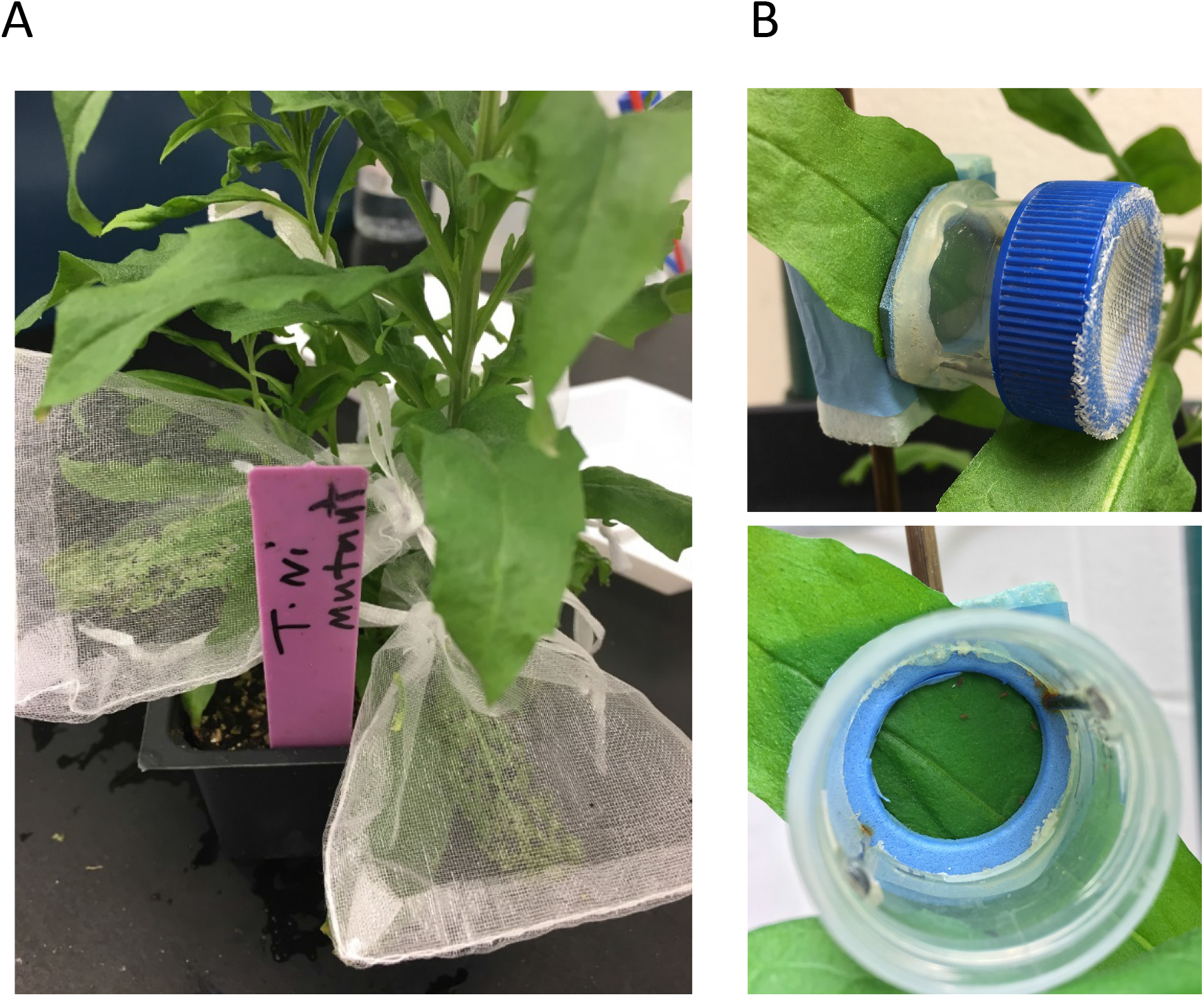
Experimental setup for insect bioassays. (A) Larval growth assays with *Trichoplusia ni*. Larvae were caged individually on leaves of *Erysimum cheiranthoides* variety Elbtalaue and the 454 mutant line. (B) *Myzus persicae* reproduction assays. Aphids were caged in groups of 5 on leaves of *E. cheiranthoides* variety Elbtalaue and the 454 mutant line.

**Fig. S2.**
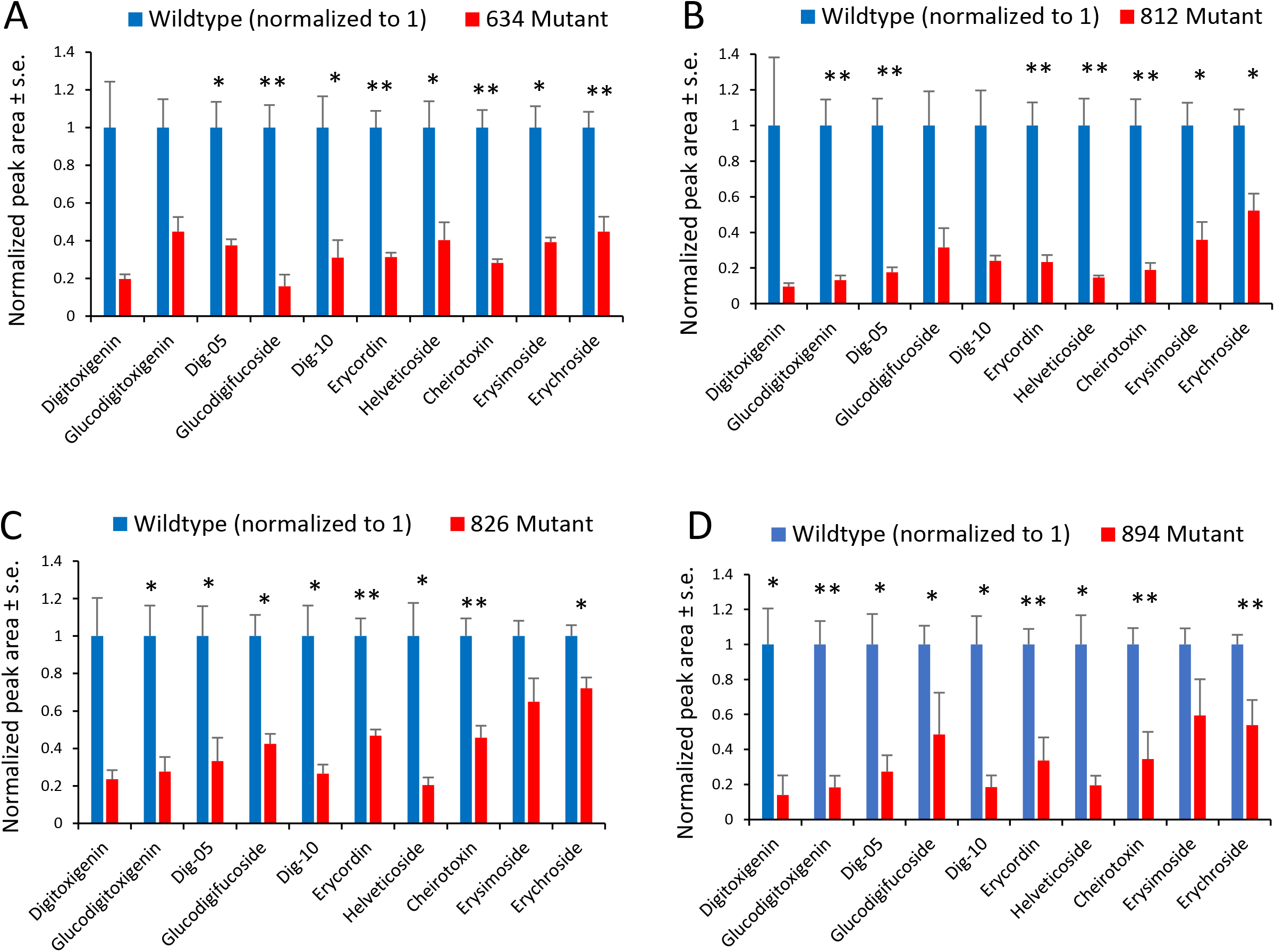
Mutant lines 634 (A), 812 (B), 826 (C), and 894 (D) have decreases in overall cardenolide content, as measured by UPLC-MS. Wildtype peak area is normalized to 1 for each cardenolide. Mean +/-s.e. of N = 5, *P < 0.05, **P < 0.01, t-test.

**Fig S3.**
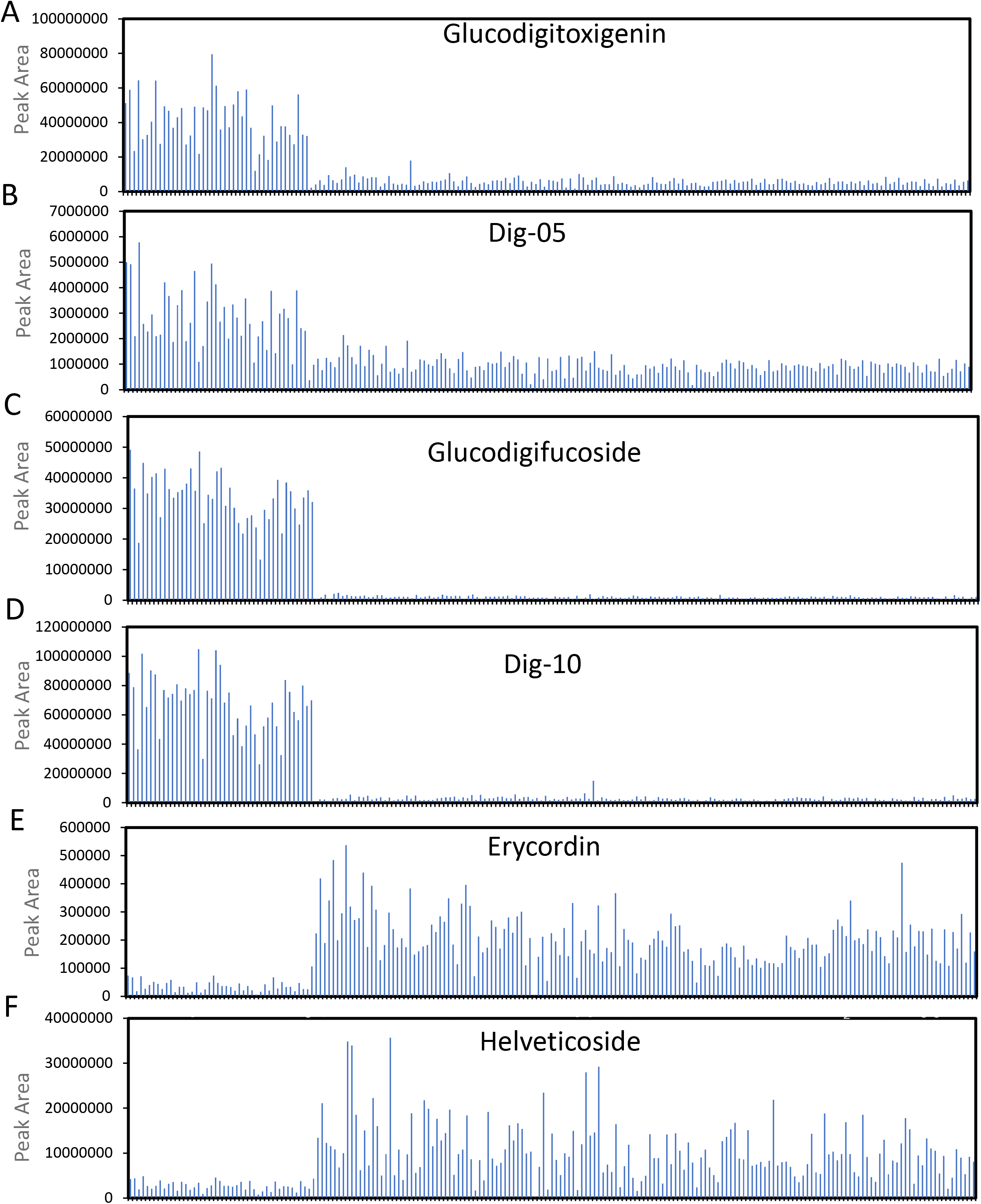
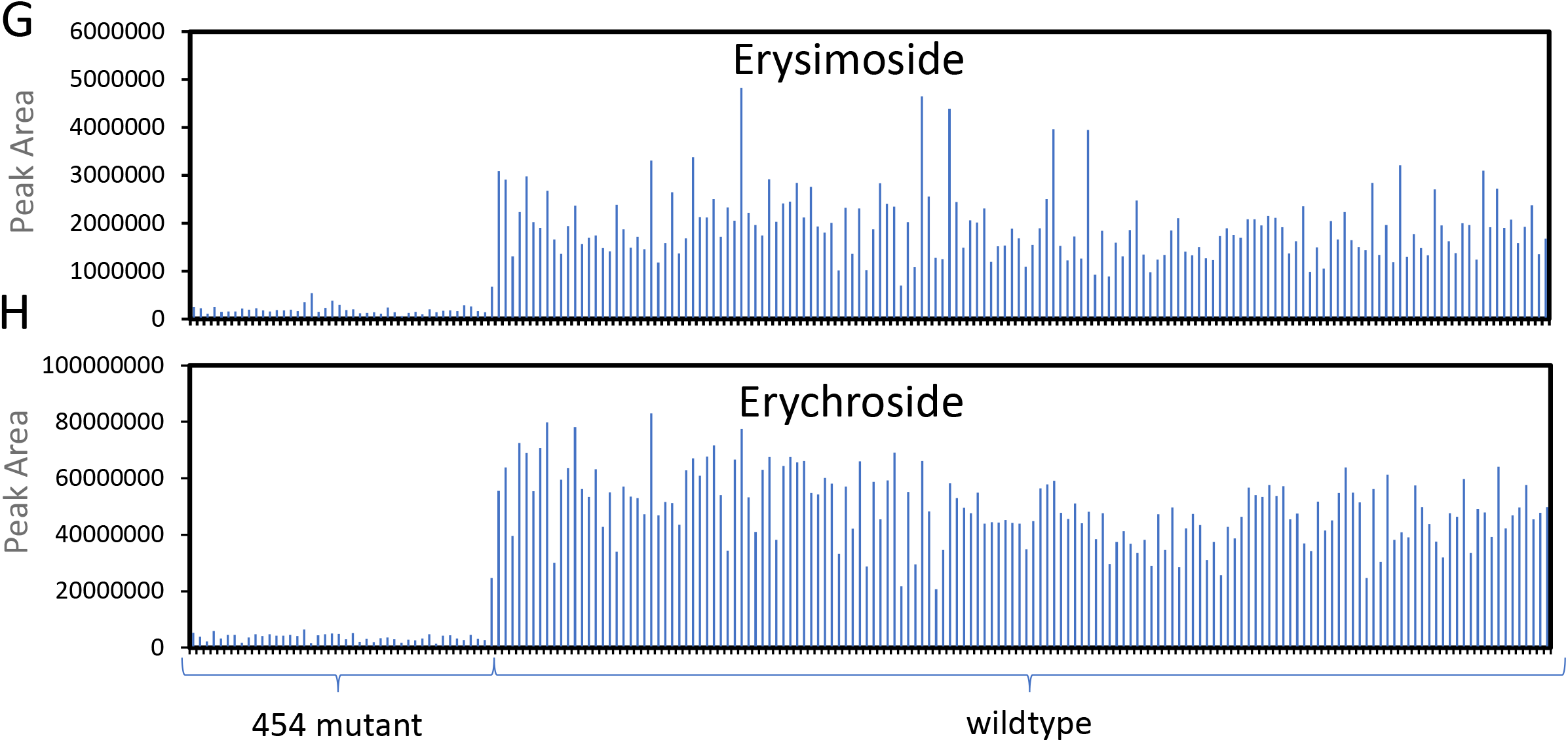
UPLC-MS analysis of 203 F2 progeny from a cross between the 454 mutant line and wildtype Elbtalaue was used to classify each plant as 454 mutant or wildtype based on the abundance of (A) glucodigitoxigenin, (B) dig-5, (C) glucodigifucoside, (D) dig-10, (E) erycordin, (F) helveticoside, (G) erysimoside, and (H) erychroside. The F2 plant samples are in the same order in each graph and have been sorted by mutant and wildtype phenotype.

## Notes

### Competing Interest Statement

The authors have declared no competing interest.

